# Naive human B cells engage the receptor binding domain of SARS-CoV-2, variants of concern, and related sarbecoviruses

**DOI:** 10.1101/2021.02.02.429458

**Authors:** Jared Feldman, Julia Bals, Clara G. Altomare, Kerri St. Denis, Evan C. Lam, Blake M. Hauser, Larance Ronsard, Maya Sangesland, Thalia Bracamonte Moreno, Vintus Okonkwo, Nathania Hartojo, Alejandro B. Balazs, Goran Bajic, Daniel Lingwood, Aaron G. Schmidt

## Abstract

Exposure to a pathogen elicits an adaptive immune response aimed to control and eradicate. Interrogating the abundance and specificity of the naive B cell repertoire contributes to understanding how to potentially elicit protective responses. Here, we isolated naive B cells from 8 seronegative human donors targeting the SARS-CoV-2 receptor-binding domain (RBD). Single B cell analysis showed diverse gene usage with no restricted complementarity determining region lengths. We show that recombinant antibodies engage SARS-CoV-2 RBD, circulating variants, and pre-emergent coronaviruses. Representative antibodies signal in a B cell activation assay and can be affinity matured through directed evolution. Structural analysis of a naive antibody in complex with spike shows a conserved mode of recognition shared with infection-induced antibodies. Lastly, both naive and affinity-matured antibodies can neutralize SARS-CoV-2. Understanding the naive repertoire may inform potential responses recognizing variants or emerging coronaviruses enabling the development of pan-coronavirus vaccines aimed at engaging germline responses.

**One Sentence Summary:** Isolation of antibody germline precursors targeting the receptor binding domain of coronaviruses.

## MAIN TEXT

Initial exposure to viral antigens by natural infection or vaccination primes an immune response and often establishes an immune memory which can prevent or control future infections. The naive repertoire contains potential B cell receptor (BCR) rearrangements capable of recognizing these antigens, often the surface-exposed glycoproteins. An early step in generating humoral immunity involves activation of these naive B cells through recognition of a cognate antigen (*1*) which in turn can lead to affinity maturation through somatic hypermutation (SHM) and subsequent differentiation (*2*). The initial engagement of the naive repertoire begins this cascade and often coincides with the eventual generation of a protective or neutralizing antibody response (*3, 4*).

For SARS-CoV-2, the etiological agent of COVID-19, the development of a neutralizing antibody response after primary infection or vaccination is associated with protection against reinfection in non-human primates (*5–9*). In humans, the presence of neutralizing antibodies can predict disease severity and survival after primary SARS-CoV-2 infection (*10*) or vaccination (*11*) and correlates with protection from symptomatic secondary infection (*12, 13*). Further, the two arms of humoral immune memory, long-lived bone marrow plasma cells (*14*) and circulating memory B cells (*15–19*), were induced by natural infection in humans and may persist for at least 8 months after primary infection providing potentially durable long-term protection. Comparable levels of neutralizing antibody titers were present in convalescent COVID-19 subjects and vaccine recipients (*20–22*) further supporting the role of adaptive immune responses in helping to control and prevent disease severity.

Both infection- and vaccine-elicited antibodies target the major envelope glycoprotein, spike, present on the virion surface (*23*). A substantial component of the neutralizing response engages the receptor binding domain (RBD) (*24–29*) and does so by directly blocking interactions with the viral receptor ACE2 (*30–35*). Isolated RBD-directed monoclonal antibodies derive from diverse heavy- and light-chain variable gene segments suggesting that multiple biochemical solutions for developing RBD-directed antibodies are encoded within the human B-cell repertoire (*24, 26, 29, 36*). Potential immunogenicity of this antigenic site is based on the human naive B cell repertoire, and the overall frequency of naive BCRs that have some level of intrinsic affinity to stimulate their elicitation (*37–40*). However, antigen-specificity of naive B cells is largely undefined.

Traditional approaches for studying antigen-specific naive B cells include bioinformatic mining of available BCR datasets and inference of likely germline precursors by “germline-reverting” mature BCR sequences, which can be limited by the availability of heavy and light chain paired sequence data and unreliable CDR3 (complementarity-determining region 3) loop approximation, respectively. Here, we address this limitation by characterizing human naive B cells specific for the SARS-CoV-2 RBD directly from the peripheral blood of seronegative donors to understand their relative abundance, intrinsic affinity, and potential for activation. Furthermore, we asked whether the SARS-CoV-2 specific naive repertoire could also engage related circulating variants of concern and pre-pandemic CoVs. We find that SARS-CoV-2 RBD-specific naive B cells were of unrestricted gene usage and several isolated B cells had affinity for circulating SARS-CoV-2 variants and related CoV-RBDs. We determined the structure of a representative naive antibody that binds the SARS-CoV-2 RBD with a mode of recognition similar to a multi-donor class of antibodies prevalent in human responses to SARS-CoV-2 infection (*41*). Further, we improved the affinity for two representative naive antibodies to RBD and showed that the starting naive specificity dictated the breadth of evolved clones to circulating variants. The analysis of the human naive antigen-specific B cell repertoire for the SARS-CoV-2 RBD and its capacity to recognize related variants and emerging CoVs may inform the rational design of epitope-focused immunogens for next generation vaccines.

### Isolated SARS-CoV-2-specific naive B cells are genetically diverse

To measure the reactivity of naive human B cells specific for the SARS-CoV-2 RBD we adapted an *ex vivo* B cell profiling approach used previously to study epitope-specific naive precursors targeting neutralizing sites on HIV (*42–44*) and influenza virus surface glycoproteins (*37*). We first designed a SARS-CoV-2 RBD construct that positions two glycans at residues 475 and 501 to selectively block binding to ACE2 and the receptor-binding motif (RBM)-directed antibody, B38 (**fig. S1**) (*45*). Using this “ΔRBM” probe, in addition to wildtype SARS-CoV-2 spike, and RBD probes, we isolated naive (CD19^+^/IgD^+^/IgG^−^) B cells specific to the RBD and, more finely, the RBM from the peripheral blood of 8 SARS-CoV-2 seronegative human donors (**Fig. 1A** and **fig. S1E**). We defined RBM-specificity as B cells that bound to fluorescently labeled spike and RBD, but not the ΔRBM probe (**fig. S2A**). Although rare, all 8 donors had detectable populations of RBM-specific naive B cells (**fig. S2B**). The median frequency of RBM-specific B cells among total and naive B cells was 0.0021% and 0.0023%, respectively (**Fig. 1B**). Within spike-reactive, naive cells, the median frequency of RBM-specific B cells was 3.2% (**Fig. 1C**); this potentially suggests that a large proportion of spike epitopes targeted by naive responses reside outside of the RBD. The majority of IgD^+^ RBM-specific B cells were CD27^−^ (mean frequency ∼97%), in agreement with the naive B cell phenotype (**fig. S2C**).

**Fig. 1.**
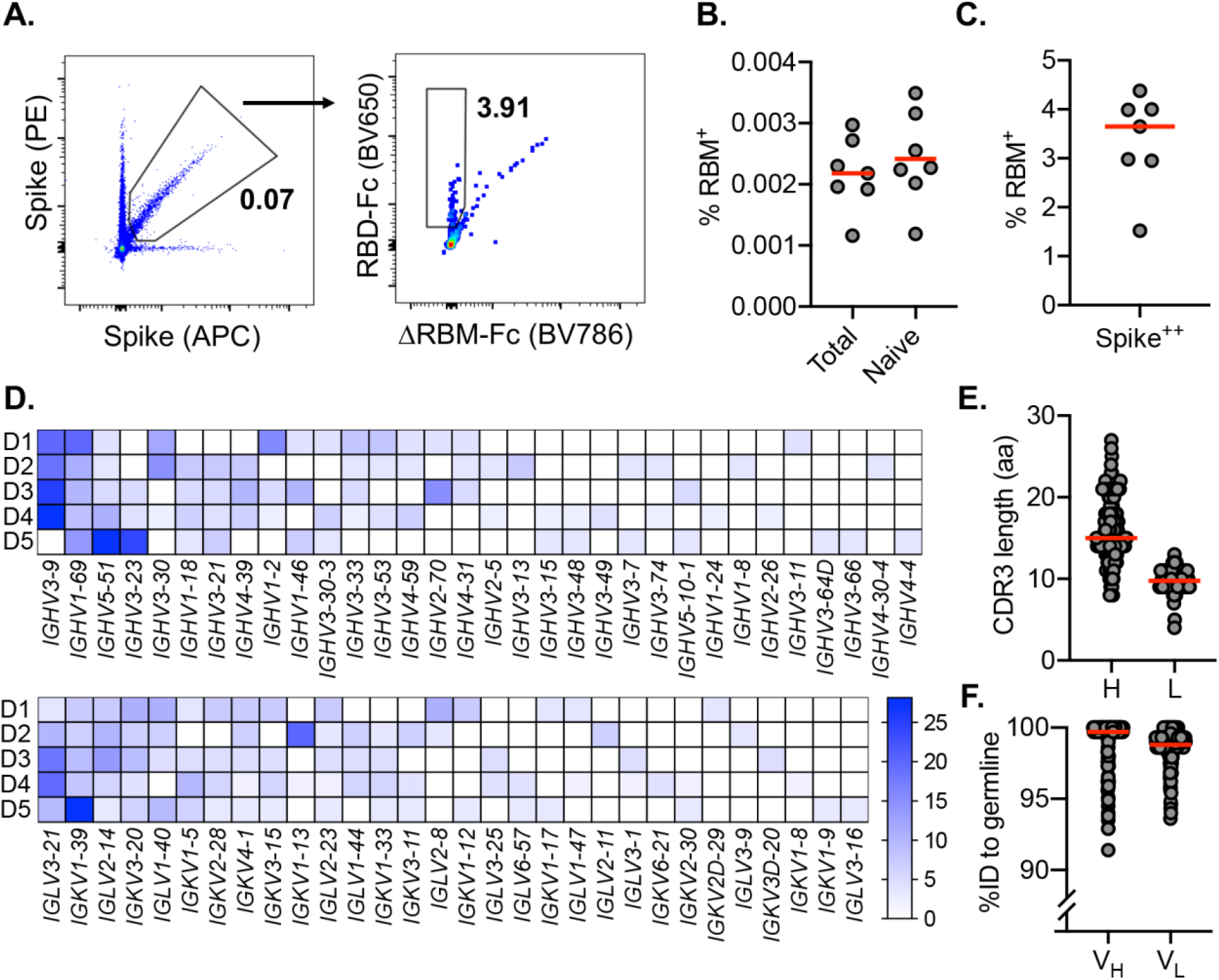
SARS-CoV-2-specific naive B cells isolation and characterization. **(A)** RBM-specific naive B cells from seronegative human donors were isolated by fluorescence-activated cell sorting gated on CD19^+^IgD^+^IgG^−^; representative plot from donors 1 and 2 is shown. ΔRBM is a sorting probe with N-linked glycans at residues 501 and 475. RBM-specific B cell frequency among (**B**) total, naive, and (**C**) spike-positive cells from each donor (*n* = 8). (**D**) Heat map showing variable-gene usage for all paired B cell sequences. Scale indicates percent of total sequences for each donor separately. (**E**) Heavy (H) and light (L) CDR3 amino acid length distribution determined using IMGT numbering. Red bars indicate median amino acid length. (**F**) Divergence from inferred germline gene sequences. Red bars indicate the median percent values.

To understand in more detail the properties of this naive repertoire, we obtained 163 paired heavy- and light-chain antibody sequences from 5 of the 8 donors **(Fig. 1D** and **Table S1)**. Sequence analysis showed that all clones were unique with diverse gene usage for both heavy and light chains and minimal gene pairing preferences (**Table S1**). These data reflect the polyclonal gene usage observed in RBD-specific memory B cells sequenced from COVID-19 convalescent individuals (*26, 29, 36*) and vaccine recipients (*23*), suggesting that a diverse pool of antibody precursors can be activated upon antigen exposure. In comparing this naive repertoire to gene usage distribution from non-SARS-CoV-2-specific repertoires (*46*), we observed an increase in mean repertoire frequency of ∼20% for IGHV3-9 in 4 out of 5 sequenced donors (**fig. S3A**). Notably, this enrichment of IGHV3-9 was also observed in isolated memory B cells from convalescent individuals (*47*) and vaccine recipients (*23*), as well as in expanded IgG^+^ B cells sequenced from a cohort of COVID-19 subjects during acute infection (*36*). These expanded clones detected shortly after symptom onset displayed low levels of SHM (*36*), suggesting potential IGHV3-9 usage in an early extrafollicular response in which naive B cells differentiate into short-lived plasma cells (*48*). Additionally, IGHV3-53 and 3-30 gene segments, over-represented in RBD-specific antibodies isolated from convalescent subjects (*27, 35, 49*), were recovered from three sequenced donors (13 total clones; ∼8.0% of total). The amino acid length of heavy and light chain third complementarity-determining regions (CDR3) ranged from 8 to 27 (average length ∼16) for HCDR3 and 4 to 13 (average length ∼10) for LCDR3 (**Fig. 1E**). These lengths are normally distributed relative to both unselected human repertoires (*46, 50*) and RBD-specific memory B cell repertoires (*23, 26, 27, 29*); this is in contrast to antibody precursors targeting the influenza and HIV receptor binding sites which have strict requirements for length (*51*) or gene usage (*52, 53*). These data suggest that overall HCDR3 length does not restrict precursor frequency and there appears no inherent bias for CDR3 length conferring RBM-specificity. The majority of obtained sequences were at germline in both the variable heavy (V_H_) and light (V_L_) chains. However, despite sorting B cells with a naive phenotype, some sequences were recovered that deviated from germline. Specifically, the V_H_ ranged from 91.4 to 100% identity to germline, with a median of 99.7%; the V_L_ ranged from 93.6 to 100%, with a median of 99.3% (**Fig. 1F, fig. 2B, C**).

**Fig. 2.**
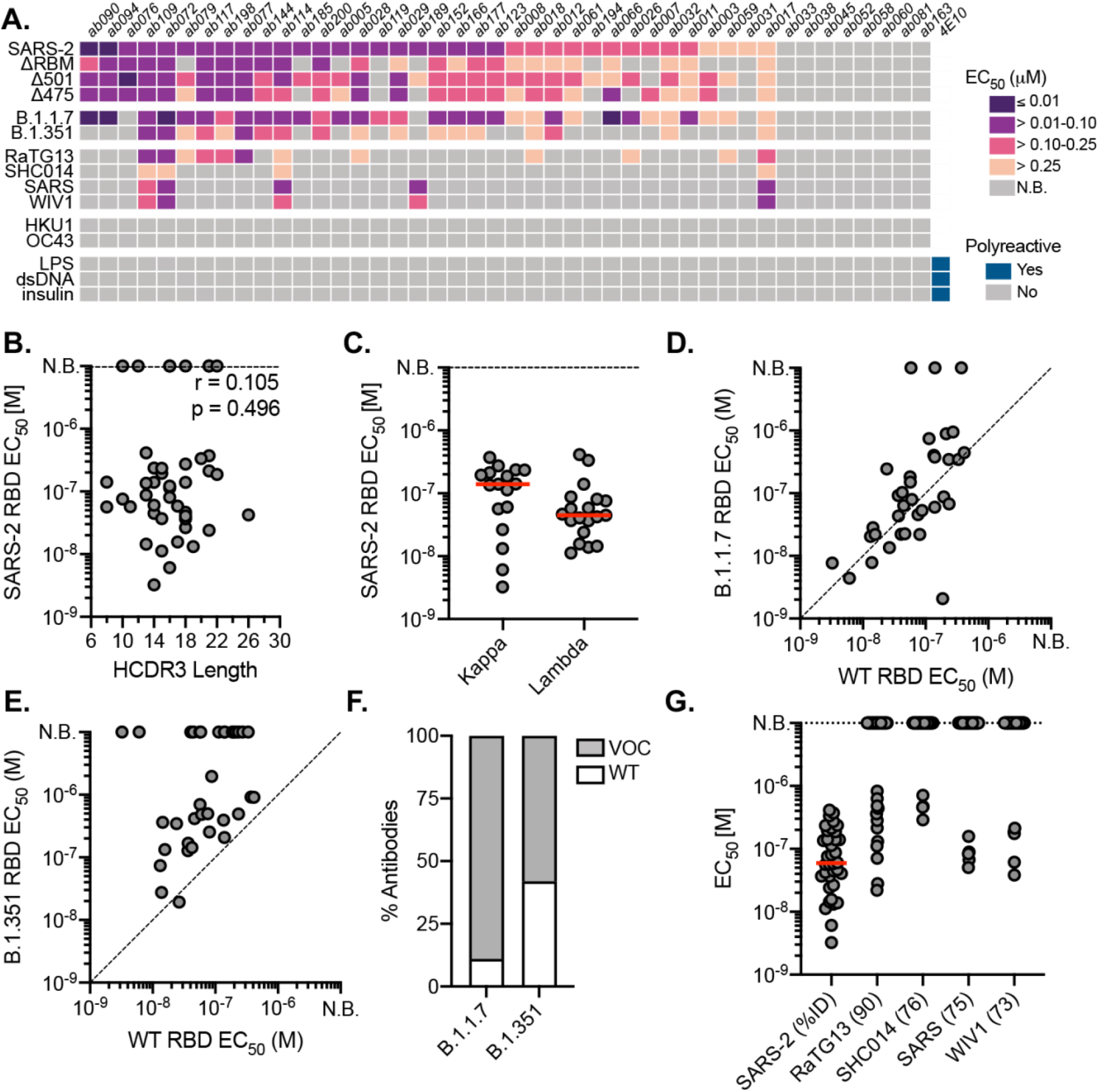
Binding properties and specificity of isolated naive antibodies. (**A**) ELISA binding heat map of 44 naive IgGs. Binding to wildtype SARS-CoV-2 RBD (SARS-2), ΔRBM, individual RBD glycan variants, circulating variants, related CoVs, hCoVs and polyreactivity antigens. (**B**) Pearson correlation analysis of SARS-CoV-2 RBD affinities and HCDR3 length. (**C**) ELISA EC_50_s for IgGs with detectable SARS-CoV-2 RBD binding (*n* = 36) based on kappa or lambda gene usage. Red bars indicate the mean EC_50_ values. (**D**) Wildtype SARS-CoV-2 RBD ELISA EC_50_s plotted against EC_50_s for B.1.1.7 RBD (**E**) B.1.351 RBD. (**F**) Proportion of SARS-CoV-2 RBD binders with detectable ELISA affinity for variants of concern (VOC) B.1.1.7 and B.1.351 RBDs. (**G**) ELISA EC_50_ values to related sarbecovirus RBDs displayed in decreasing order of paired-sequence identity.

### Naive antibodies engage SARS-CoV-2 RBD with high affinity

To obtain affinities of the isolated naive antibodies, we cloned and recombinantly expressed 44 IgGs selected to reflect the polyclonal RBD-specific repertoire with representatives from diverse variable region gene segments (**Table S1**). Additionally, we ensured diversity in terms of HCDR3 length, kappa and lambda usage, as well as representation from all 5 donors. By ELISA, we identified IgGs with detectable binding to SARS-CoV-2 RBD; we summarize these results for all antibodies (**Fig. 2A**) and parsed by donor (**fig. S3D**). Across 5 donors, 36 (∼81%) bound to monomeric SARS-CoV-2 RBD (**Fig. 2A**) with EC_50_ values ranging from 3.3 to 410 nM and a mean of 62 nM (**Fig. 2A** and **fig. S3E**). These antibodies included 32 unique variable heavy and light chain pairings (**Table S1**). Of the binding population, there is no apparent predisposition for HCDR3 length or light chain pairing (**Fig. 2 C, D**). We further defined the epitopic region of these IgGs using the ΔRBM construct and the individual glycan variants, Δ501 and Δ475, both of which independently block ACE2 cell-surface binding but are on opposite sides of the RBM (**fig. S1E, F**). 11 IgGs had no detectable ΔRBM binding (e.g., ab079, ab119), while 21 IgGs had reduced ELISA binding relative to wild-type RBD, reflected in the reduced ΔRBM median EC_50_ values (**fig. S3E**). We also identified examples of antibodies sensitive to only Δ475 (e.g., ab185) and only Δ501 (e.g., ab007) (**Fig. 2A** and **fig. S3E).**

To obtain binding kinetics independent of avidity effects from bivalent IgGs, 12 antibodies were selected for expression as Fabs to determine monovalent binding affinity (*K_D_s*) by biolayer interferometry (BLI). Using monomeric RBD as the analyte, 10 of the 12 Fabs had detectable binding with *K_D_s* ranging from ∼6.5 to ∼75 µM; the other two remaining Fabs (ab177, ab185), gave unreliable affinity measurements (i.e., >100 µM) (**fig. S4**). Notably, all Fabs had characteristically fast off rates (*k_off_)*. This observation is consistent for germline B cells where fast off-rates are compensated by avidity due to overall BCR surface density (*54*); subsequent affinity gains via SHM often result in slowing of the off-rate and is a canonical mechanism of improved antigen binding (*55–57*).

**Fig. 4.**
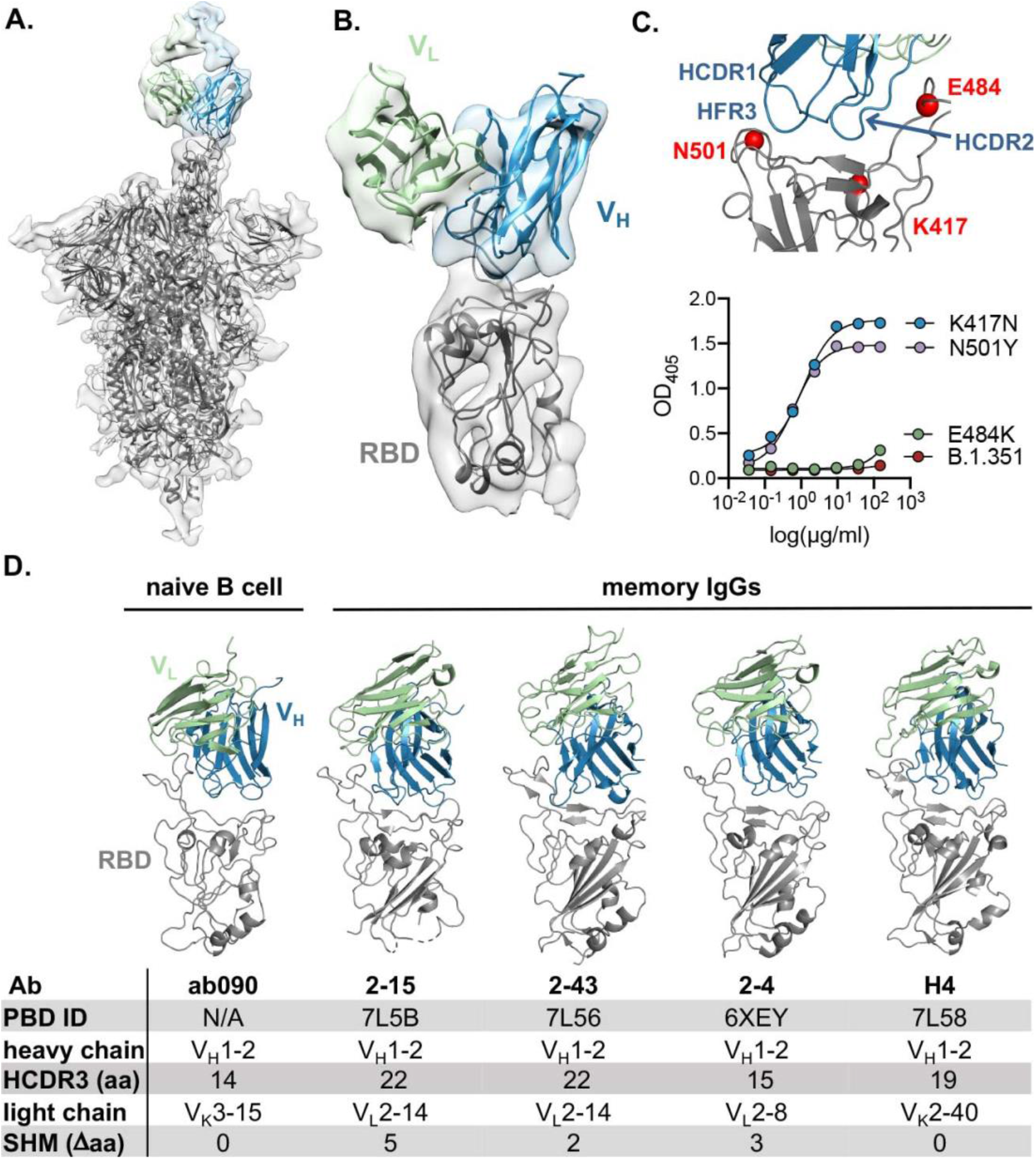
ab090 recognizes the SARS-CoV-2 RBM. (**A**) Cryo-EM structure of the SARS-CoV-2 spike trimer (grey) with ab090 Fab bound to one RBD in the up position. (**B**) ab090 recognizes the SARS-CoV-2 RBM with through a paratope centered on the V_H_ (blue). (**C**) Close-up view showing the approximate locations of HCDR loops proximal to the RBM epitope and B.1.351 RBD mutations highlighted in red (top). ELISA binding reactivity of ab090 to individual mutations from B.1.351 RBD (bottom). (**D**) ab090 binds to the RBM with a similar mode and angle of approach to IGHV1-2 neutralizing antibodies isolated from memory B cells from convalescent COVID-19 donors. RBDs (grey) are shown in the same relative orientation in each panel with PBD codes and sequence attributes listed in the below.

### Naive antibodies engage SARS-CoV-2 variants of concern

The emergence of SARS-CoV-2 variants with mutations in RBD has raised significant concern that antigenic evolution will impair recognition of RBD-directed antibodies elicited by prior infection and vaccination with an antigenically distinct SARS-CoV-2 variant (*58–61*). We therefore asked whether these naive antibodies, isolated using wild-type SARS-CoV-2 RBD, could recognize circulating viral variants, B.1.1.7 (mutations N501Y) (*62*) and B.1.351 (mutations K417N/E484K/N501Y) (*63*); the former has now become the most prevalent circulating variant in the US and many other countries (*64*). we find that 89% of the antibodies with wild-type RBD affinity also bound to the B.1.1.7 variant with a comparable mean affinity of 68.9 nM (**Fig. 2D, F**). For B.1.351, a concerning variant prevalent in South Africa (*64*), 62% of the wild-type SARS-CoV-2 RBD binding IgGs also bound to the B.1.351 variant, many of which displayed reduced ELISA binding relative to wild-type RBD with a mean affinity of 262 nM (**Fig. 2E, F**). A more pronounced reduction in cross-reactivity to the B.1.351 variant may be predictive of reduced sera binding and neutralization titers from convalescent individuals and vaccine-recipients (*22, 58, 65*).

### Naive antibodies engage pre-emerging CoVs

We next tested the cross-reactivity of these naive antibodies to related sarbecovirus RBDs, which also use ACE2 as a host receptor (*66*). Our panel included the previously circulating SARS-CoV RBD and representative preemergent bat CoV RBDs from WIV1 (*67*), RaTG13 (*68*), and SHC014 (*69*). These RBDs share 73 to 90% paired-sequence identity with the highest degree of amino acid conservation in residues outside of the RBM (*70*). 13 antibodies cross-reacted with at least one additional RBD in our panel, with decreasing affinity for RBDs with more divergent amino acid sequence identity (**Fig. 2A, G**). Notably, ab017, ab072, ab109, and ab114 had broad reactivity to all tested sarbecovirus RBDs, suggesting binding to highly conserved epitopes. Of these cross-reactive antibodies, ab017 and ab114, derive from the same IGHV3-33 and IGVL2-14 paring but were isolated from different donors, suggesting a shared or public clonotype.

### Naive antibodies are not polyreactive and do not engage seasonal coronaviruses

Prior studies have shown that germline antibodies are more likely to display polyreactivity relative to affinity-matured antibodies with higher levels of SHM from mature B cell compartments (*71–74*). We therefore tested the polyreactivity of all 44 naive antibodies using three common autoantigens, double-stranded DNA (dsDNA), *Escherichia coli* lipopolysaccharide (LPS), and human insulin in ELISA (**Fig. 2A**) We observed no polyreactivity of any naive antibody, including those that are broadly reactive. Furthermore, none of the naive antibodies bound RBDs from the human seasonal betacoronaviruses (hCoVs), OC43 and HKU1 (**Fig. 2A**), which share 22 and 19% paired-sequence identity to SARS-CoV-2 RBD, respectively. Together, these results suggest that the isolated naive B cells encode BCRs with specificity to sarbecoviruses.

### *In vitro* reconstitution of naive B cell activation

Physiological interactions between a naive BCR and cognate antigen occurs at the B cell surface. Naive BCRs are displayed as a bivalent membrane-bound IgM and multivalent antigen binding can initiate intracellular signaling resulting in an activated B cell with the capacity to differentiate to antibody secreting plasma cells or memory cells (*75*). To determine whether the isolated RBD-specific naive BCRs have the capacity to be activated, we generated stable Ramos B cell lines expressing ab090 or ab072 as cell-surface BCRs and measured their activation by monitoring calcium flux *in vitro* (*76*). These antibodies were selected to represent divergent germline gene usage and specificities: 1) ab090 (IGHV1-2/IGKV3-15) bound SARS-CoV-2 and variant B.1.1.7 RBDs, but not variant B.1.351 and WIV1 RBDs (**Fig. 3A**); and 2) ab072 (IGHV3-23/IGLV2-14) had broad reactivity to all RBDs (**Fig. 3B**). To assess BCR activation, we generated ferritin-based nanoparticles (NPs) for multivalent RBD display using SpyTag-SpyCatcher (*70, 77, 78*); these RBD NPs included SARS-CoV-2, B.1.1.7, B.1.351 and WIV RBDs. We found that ab090 expressing Ramos B cells were only activated by SARS-CoV-2 RBD and variant B.1.1.7 RBD NPs (**Fig. 3C**), while ab072 Ramos B cells were activated by all RBD-NPs (**Fig. 3D**). Notably, these data parallel the observed recombinant binding specificity of each antibody. Importantly, neither ab090 nor ab072 Ramos B cell lines were activated by influenza hemagglutinin NPs, suggesting that this activation is sarbecovirus RBD-specific (**Fig. 3C, D**).

**Fig. 3.**
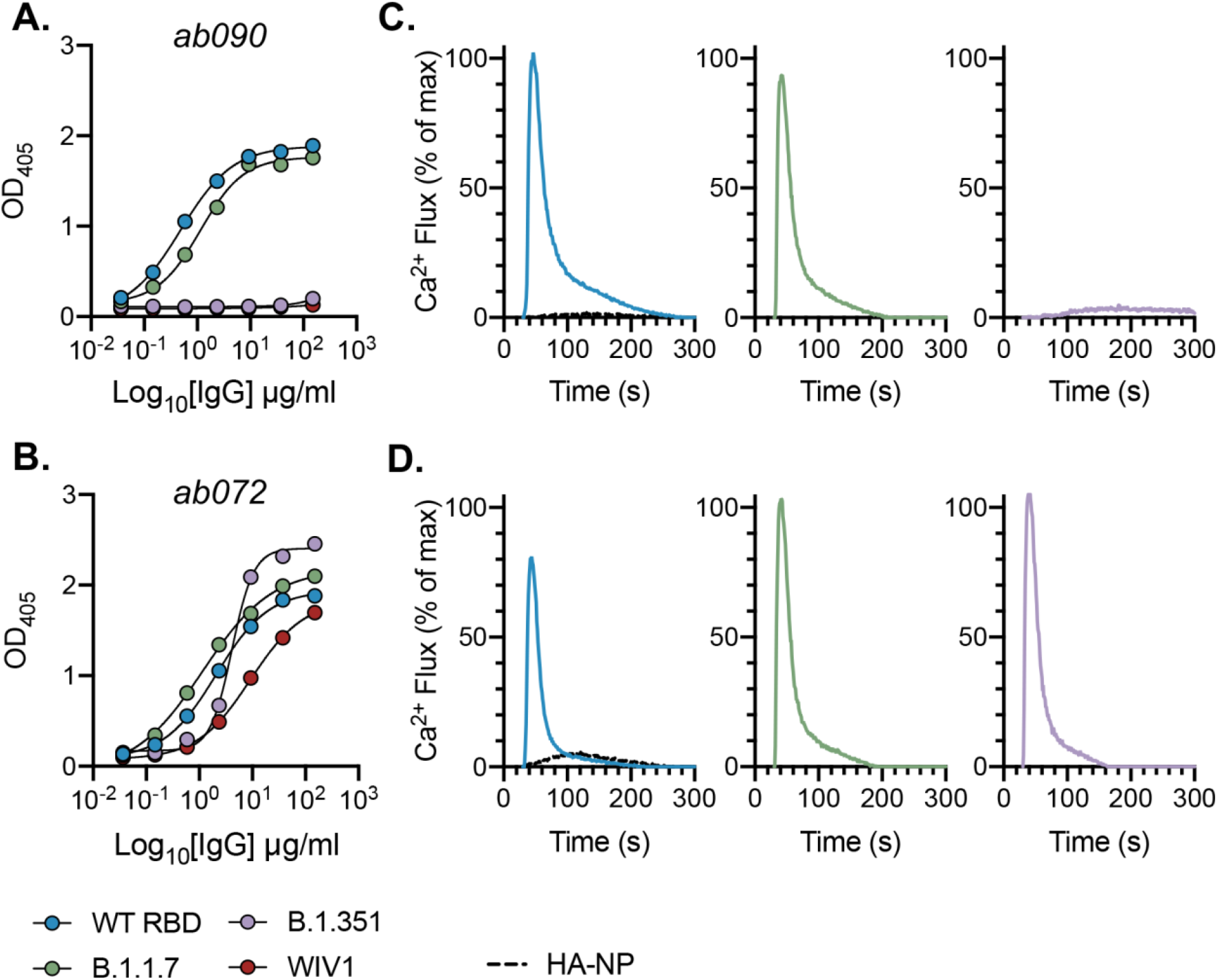
*In vitro* reconstitution of naive B cell activation. (**A**) ELISA binding reactivity shows restricted specificity of ab090 and (**B**) broad binding of ab072 to wildtype (WT) SARS-CoV-2, B.1.1.7, B.1.351, and WIV1 RBDs. (**C**) BCR activation as measured by calcium flux in a Ramos B cell line expressing ab090 membrane-anchored IgM (mIgM) and (**D**) and ab072 mIgM in response to ferritin nanoparticles (NPs) displaying WT SARS-CoV-2, B.1.1.7, B.1.351, and WIV1 RBDs. Influenza hemagglutinin (HA) NP was used as a negative control.

### ab090 engages the SARS-CoV-2 RBM

To further characterize the epitope specificity of a representative naive antibody, we determined the structure of ab090 in complex with SARS-CoV-2 spike (S) by electron cryomicroscopy (cryo-EM). A ∼6.7-Å structure showed one Fab bound to an RBD in the “up” conformation (**Fig. 4A, B and fig. S5)**. Based on this modest resolution structure, we make the following general descriptions of the antibody-antigen interface. The interaction between ab090 and the RBD is mediated primarily by the antibody heavy chain, with the germline encoded HCDR1, HCDR2, and the framework 3 DE-loop centered over the RBM epitope (**Fig. 4B**). The ab090 light chain is oriented distal to the RBD and does not appear to substantially contribute to the paratope (**Fig. 4B**). IGHV1-2 antibodies represent a prevalent antibody class in human responses to SARS-CoV-2 infection, many of which display high neutralization potency (*41, 79*). ab090 shares a V_H_-centric mode of contact and angle of approach similar to members of this class of infection-elicited antibodies (**Fig. 4D**), despite varying HCDR3 lengths and diverse light chain pairings (**Fig. 4D**) (*41, 79*). Additionally, members of the IGHV1-2 antibody class contain relatively few SHMs (**fig. S5B**). We note that many of the infection-elicited IGHV1-2 RBD-specific memory B cells derive from the IGHV1-2*02 allele, while ab090 is encoded by the IGHV1-2*06 allelic variant (**fig. S5B**). The IGHV1-2*06 allele is represented by a single nucleotide polymorphism encoding an arginine rather than a tryptophan at position 50 (*80*) (**fig. S5B**). Notably, a potent neutralizing antibody, H4, derives from the same *06 allele (*34*). In conjunction with the structure, we biochemically defined the sensitivity of ab090 to variant B.1.351 by testing the binding to individual mutations. Binding affinity was detected to SARS-CoV-2 RBDs with either N501Y or K417N mutations, but not to E484K alone (**Fig. 4C**). Based on the structure, the E484K mutation, is grossly positioned proximal to the CDRH2 loop (**Fig. 4C**), which has a germline-encoded motif critical for IGHV1-2 antibody binding to RBD (**fig. S5B**) (*41*). Indeed, infection-elicited IGHV1-2 antibodies are susceptible to escape by E484K alone, which disrupts a CDRH2 hydrogen binding network (*81*). Together, the cryo-EM structure and binding data suggest that ab090 represents a precursor of a class of RBM directed SARS-CoV-2 neutralizing antibodies. More generally, structural characterization of germline antibody complexes has been limited to hapten antigens (*82*), simple peptides (*83*) and to protein antigens bearing engineered affinity to inferred germline sequences(*84*). We present, to our knowledge, the first structure of a naturally occurring naive human antibody bound to non-engineered viral protein.

**Fig. 5.**
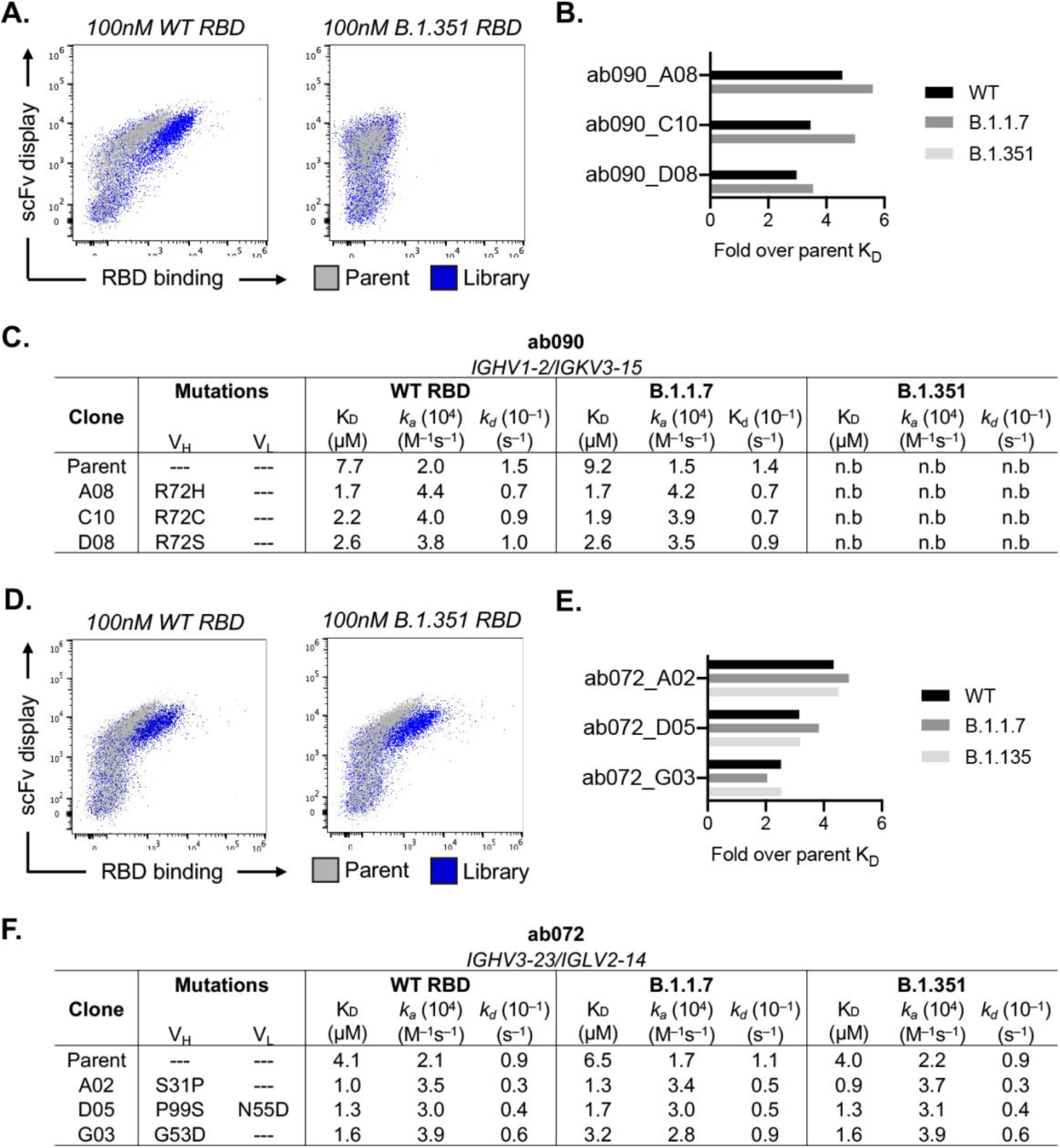
*in vitro* affinity-matured naive antibodies retain intrinsic specificity. (**A**) Enrichment of ab090 parent (grey) and affinity matured (blue) libraires to 100nM SARS-CoV-2 or B.1.351 RBD using flow cytometry (**B**) Fold enrichment in monovalent *K_D_* over ab090 parent for selected affinity matured progeny. (**C**) Kinetic analysis using biolayer interferometry (BLI) for ab090 parent and progeny Fabs to monomeric WT and variant RBDs. (**D**) Enrichment of ab072 parent (grey) and affinity matured (blue) libraires to 100nM SARS-CoV-2 or B.1.351 RBD using flow cytometry (**E**) Fold enrichment in monovalent *K_D_* over ab072 parent for selected affinity matured progeny. (**F**) Kinetic analysis using biolayer interferometry (BLI) for ab072 parent and progeny Fabs to monomeric WT and variant RBDs.

### *in vitro* affinity-matured naive antibodies retain intrinsic specificity

After initial antigen recognition and subsequent activation, naive B cells can undergo successive rounds of somatic hypermutation within the germinal center (GC) that ultimately result in higher affinity antibodies for the cognate antigen. To determine how somatic hypermutation might influence overall affinity and specificity, we used yeast surface display to *in vitro* mature ab072 and ab090. We randomly mutagenized the single chain variable fragment (scFv) variable heavy and light chain regions to generate ab072 and ab090 variant display libraries (*85*). After two rounds of selections using SARS-CoV-2 RBD, we enriched the ab072 and ab090 libraries for improved binding over their respective parental clones (**Fig. 5A, D and fig. S6A**). We also observed increased binding to B.1.351 for the ab072 library but not for ab090; notably this corresponded with the respective specificity of the parent clones (**Fig. 5A, D**).

We next isolated and sequenced individual clones from the enriched libraries. For ab090, we observed a dominant mutation, R72H, in the FRWH3 region present in 60% of sequenced clones (**fig. S6B**). Notably, multiple mutations at position 72 conferred a ∼3- to 5-fold improvement in monovalent affinity relative to parental ab090 for wild-type and B.1.1.7 RBDs, with no detectable B.1.351 binding for affinity matured progeny (**Fig. 5B, C**). We observed no mutations within the light chain which appears to be consistent with the V_H_-centric binding mode in the cryo-EM structure (**Fig. 4**). For the broadly reactive ab072, isolated clones had mutations in both the V_H_ and V_L_ ; ∼35% of the sequenced clones had mutation S31P in the HCDR1 (**fig. S6B, C**). There was 3- to ∼5-fold improvement in monovalent affinity of ab072 progeny relative to parent for SARS-CoV-2, B.1.1.7 and B.1.351 RBDs (**Fig. 5E, F**). Collectively, these data identify potential mutations that can improve affinity while retaining initial parental antigen specificity.

### SARS-CoV-2 pseudovirus neutralization by naive and affinity-matured Abs

We next used a SARS-CoV-2 pseudovirus assay (*10*) to ask whether any of the isolated naive antibodies and affinity matured clones were capable of blocking transduction of target cells. We found that of the 36 RBD-binding antibodies tested in this assay, 5 had detectable levels of neutralization (∼14%) (**Fig. 6A**). These antibodies, obtained from multiple donors, have no commonality with respect to their gene usages and HCDR3 lengths (**Fig. 6B**). While these naive antibodies were not as potent as B38, isolated from a memory B cell (*34*), the observation, nevertheless, that the naive repertoire has antibodies that neutralize is noteworthy.

**Fig. 6.**
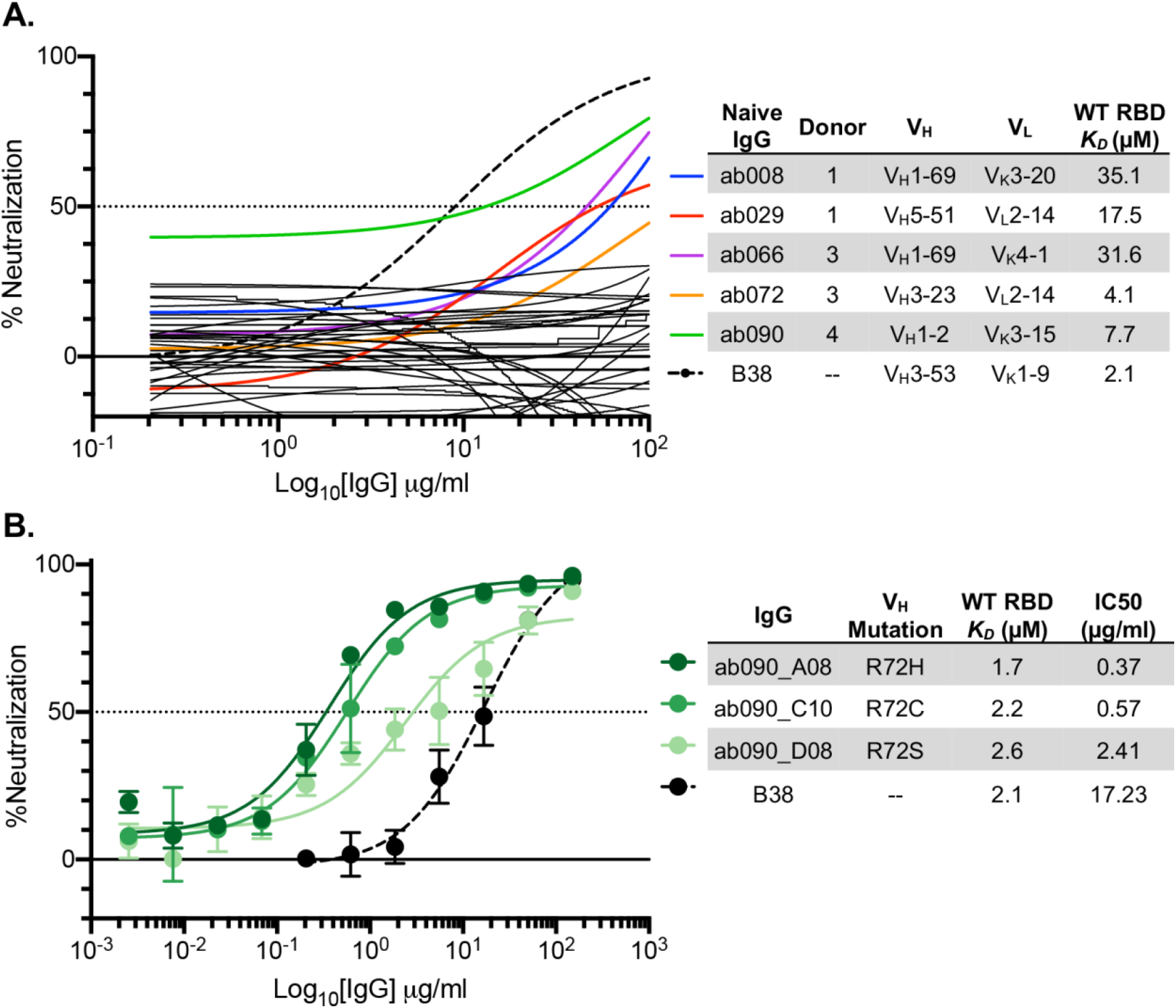
SARS-CoV-2 pseudovirus neutralization by naive and affinity-matured Abs. (**A**) SARS-CoV-2 pseudovirus neutralization assay for 36 purified IgGs. Curves in color highlighted antibodies with neutralizing activity with donor and monovalent wild-type RBD affinity listed for this subset of antibodies. The neutralizing monoclonal antibody, B38, was used as a positive control. Dashed lines indicate IC_50_ values and data represent means ± SD of two technical replicates. (**B**) SARS-CoV-2 pseudovirus neutralization for select affinity matured progeny from the ab090 lineage with respective mutations relative to ab090 parent sequence, monovalent wild-type RBD affinity, and IC50 listed.

To determine whether improved affinity correlated with enhanced neutralization potency, we evaluated the affinity matured progeny of ab090 in a SARS-CoV-2 pseudovirus neutralization assay (**Fig. 6B**). We find that all three ab090 progeny that had higher affinity for SARS-CoV-2 RBD also had increased neutralization potency. ab090_A08 bearing the R72H mutation had the highest affinity gain and was the most potent neutralizer with a *K_D_* of 1.7μM and an IC50 of 0.37 µg/ml, respectively. Notably, ab090 progeny had IC50 values similar to other IGHV1-2 memory B cells isolated from convalescent donors (*41*); this increase in potency is conferred through minimal somatic hypermutation.

## DISCUSSION

The development of a protective humoral immune response upon infection or vaccination relies on the recruitment, activation, and maturation of antigen-specific naive B cells. However, the specificity of the naive B cell repertoire remains largely undefined. Here, we showed that coronavirus-specific naive B cells are present across distinct seronegative donors, are of unrestricted gene usage and when recombinantly expressed as IgGs, have affinity for SARS-CoV-2 RBD, circulating variants of concern, and at least four related coronaviruses. These data suggest that RBD-specific precursors are likely present across a large fraction of individual human naive repertories, consistent with longitudinal studies of SARS-CoV-2 infected individuals in which most convalescent individuals seroconverted with detectable RBD serum antibodies and neutralization titers (*17, 86, 87*). The naive B cells characterized here engage epitopes across the RBM with a range of angles of approach as defined by our glycan variant probes and cross-reactivity profiles; this is also consistent with infection and vaccine elicited, RBD-specific repertoire characterized by epitope-mapping, deep mutational scanning and structural analyses (*30, 32, 88*). Having naive BCRs recognizing distinct or partially overlapping epitopes across the RBM may be advantageous for eliciting a polyclonal response more able to recognize variants of concern.

The presence of broadly reactive naive B cells inherently capable of recognizing sarbecovirus RBDs and circulating variants suggests that these precursors could be vaccine-amplified. Recent work showed that uninfected individuals have pre-existing SARS-CoV-2 S-reactive serum antibodies (*89–91*) and memory B cells (*28, 92*) which cross-react with hCoVs and can be boosted upon SARS-CoV-2 infection. These cross-reactive antibodies appear to be specific to the S2 domain and are predominantly IgG or IgA. Notably, this observation contrasts the cross-reactive B cells described here that engage the RBD, have no reactivity to hCoV and are IgG^−^ naive B cells suggesting that they are distinct from previously described S-reactive pre-existing antibodies.

Data suggests that the competitive success of a naive B cell within a GC is influenced by precursor frequencies and antigen affinities (*40*). However, the biologically relevant affinities necessary for activation remain unclear—indeed several studies suggest that B cell activation and affinity maturation is not restricted by immeasurably low affinity BCR interactions (*93–95*). Recently, two studies involving naive precursors of receptor-binding site (RBS) directed HIV-1 broadly neutralizing antibodies (bnAbs) contributed to our understanding of these parameters (*38, 39*). Using an *in vivo* murine adoptive transfer model, these RBS-directed precursors were recruited into a GC reaction at a precursor frequency of ∼1:10,000 and a monovalent antigen affinity of 14µM (*39*). For comparison, here we defined the SARS CoV-2 RBM-specific naive precursor frequency as 1:41,000 by flow cytometric gating (**fig. S2**) with monovalent affinities ranging from 6.5 to >100µM. These data suggest that these isolated naive B cells, especially those with demonstrable monomeric affinity, could be readily elicited upon antigen exposure. However, longitudinal studies tracking antigen specific naive B cells pre- and post-exposure are required to determine the fate (i.e., plasma cell, memory, or germinal center B cell compartments) of potential precursors and define relevant naive affinities for elicitation by SARS-CoV-2.

Through biochemical and structural analyses, we characterized a naive antibody, ab090, which resembles a commonly elicited class of potent neutralizing antibodies utilizing the IGHV1-2 gene (*41*). This class of antibodies share restricted binding specificity for wild-type SARS-CoV RBD (the vaccine strain) and the prevalent B.1.1.7 variant. This recombinant binding pattern also paralleled the reconstituted *in vitro* B cell activation dynamics of ab090 in the highly avid assay with the capacity to detect immeasurably low affinity interactions (*54*). *In vitro* affinity maturation of ab090 against corresponded to a single H-FR3 mutation, which improved monovalent affinity ∼5-fold to wild-type SARS-CoV-2 and B.1.1.7 RBDs relative to parent and pseudovirus neutralization to IC50 values less than 1µg/ml. This observation is consistent with the low levels of SHM within IGHV1-2 neutralizing antibodies (*41*) and with reports of other potent RBD-directed neutralizing antibodies with a limited level of somatic hypermutation (*24, 26, 29, 36, 96*). Further, a recent study monitoring RBD-specific memory B cell evolution up to 12 months after SARS-CoV-2 infection revealed examples of affinity matured clones with increased neutralizing breadth over time against circulating RBD variants (*97*). While *in vitro* affinity gains and neutralization potency are generally correlated (*98*), we note that affinity does not necessarily correlate to neutralization potency for all SARS-CoV-2 RBD targeting antibodies, where fine epitope specificity appears to be most relevant (*28, 99*).

Probing and characterizing the human naive B cell antigen-specific repertoire can identify precursors for vaccine or infection-specific naive B cells and expand our understanding of basic B cell biology. Germline-endowed specificity for neutralizing antibody targets on the RBD may also contribute to the strong clinical efficacy observed for the current SARS-CoV-2 vaccines (*100, 101*). Furthermore, understanding the naive B cell repertoire to potential pandemic coronaviruses may reveal commonalties in antigen-specific precursors, enabling the development of pan-coronavirus vaccines aimed at engaging broadly protective germline responses.

## METHODS

### Donor Samples

PBMCs were isolated from blood donors obtained from the MGH blood donor center (8 donors total). Prior to donating blood, subjects were required to sign a donor attestation/consent statement, as per hospital requirements, stating ‘‘I give permission for my blood to be used for transfusion to patients or for research’’. The gender and age are not recorded, however eligible donors are of at least 16 years old and weigh a minimum of 110lbs. All experiments were conducted with MGH Institutional Biosafety Committee approval (MGH protocol 2014B000035). Isolated PBMCs were used for B cell enrichment and single cell sorting (described below); plasma was aliquoted and stored at -80 °C until further use. Additionally, the control convalescent sera used for ELISA was obtained under the approved Partners Institutional Review Board (protocol 2020P000895) for use of patient samples for the development and validation of SARS-CoV-2 diagnostic tests (*10*).

### Expression and purification of recombinant CoV Antigens

Plasmids encoding the receptor binding domains (RBDs) were designed based on GenBank sequences MN975262.1 (SARS-CoV-2), ABD72970.1 (SARS-CoV), AGZ48828.1 (WIV-1), MN996532.2 (RaTG13), QJE50589.1 (SHC014), AAT98580.1 (HKU1), and AAT84362 (OC43). Constructs were codon optimized and synthesized by IDT. QuikChange Mutagenesis (Agilent) was used to insert glycosylation sites at SARS-CoV-2 RBD residues 501 and/or 475 as well as for RBD variant mutations, B.1.351 (K417N/E484K/N501Y) and B.1.1.7 (N501Y). SARS-CoV-2 spike contained a C-terminal foldon trimerization domain and HRV 3C-cleavable 6xHis and 2xStrep II tags (*102*). All proteins were transiently expressed in Expi293F cells (ThermoFisher). 5 to 7 days post-transfection, supernatants were harvested by centrifugation and further purified using immobilized metal affinity chromatography (IMAC) with cobalt-TALON resin (Takara) followed by Superdex 200 Increase 10/300 GL size exclusion column (GE Healthcare).

### Expression and purification IgGs and Fabs

IgG and Fab genes for the heavy- and light-chain variable domains were synthesized and codon optimized by IDT and subcloned into pVRC protein expression vectors and sequence confirmed (Genewiz). Fabs and IgGs were similarly expressed and purified as described above for RBDs. IgGs were buffer exchanged into PBS while Fabs were concentrated and further purified by Superdex 200 Increase 10/300 GL size exclusion column.

### ELISA

Both sera and monoclonal antibody reactivity to CoV antigens were assayed by ELISA. Briefly, 96-well plates (Corning) were coated with 5 µg/ml of monomeric RBDs in PBS at 100µl/well and incubated overnight at 4°C. Plates were blocked with 1% BSA in PBS containing 1% Tween-20 (PBS-T) for 1hr at room temperature (RT). Blocking solution was discarded and 4-fold serial dilutions of human plasma (1:20 starting dilution) or isolated monoclonal antibodies (150 µg/ml starting concentration) in PBS were added to wells and incubated for 1hr at RT. Plates were then washed three times with PBS-T. Secondary, anti-human IgG-HRP (Abcam), was added to each well at 1:20,000 dilution in PBS-T and incubated for 1hr at RT. Plates were then washed three times with PBS-T and developed with 1-Step ABTS substrate (ThermoFisher) per manufacturer recommendations. Absorbance was measured using a plate reader at 405nm. EC_50_ values were determined for monoclonal antibodies by non-linear regression (sigmoidal) using GraphPad Prism 8.4.3 software. ELISAs against OC43 and HKU1 RBDs were done at a single IgG concentration (150 µg/ml) in replicate. Positive binding was defined by an OD_405_ ≥ 0.30.

For polyreactivity ELISAs against human insulin (MilliporeSigma) and dsDNA (Calf Thymus DNA; Invitrogen), plates were coated with 2µg/ml and 50µg/ml, respectively, in PBS at 100µl/well and incubated overnight at 4°C. Plates were then blocked and incubated with IgGs as described above for CoV antigens. LPS ELISAs were measured according to a previously described method (*103, 104*). Briefly, plates were coated with 30µg/ml LPS (*Escherichia coli* O55:B5; MilliporeSigma) in carbonate buffer (100mM Na_2_CO_2_, 20mM EDTA, pH 9.6) at 100µl/well for 3hrs at 37°C, washed three times with water, and air-dried overnight at RT. Coated plates were blocked with 200µl/well of HS buffer (50mM HEPES, 0.15mM NaCl, pH 7.4) plus 10mg/ml. Plates were incubated with IgGs diluted in HS buffer containing 1mg/ml BSA for 3hrs at 37°C, washed three times with HS buffer, and developed as detailed above for CoV antigens. All polyreacivity ELISAs were performed at a single IgG concentration (15µg/ml) in replicate with positive binding was defined by an OD_405_ ≥ 0.30.

### ACE-2 cell binding assay

ACE-2 expressing 293T cells were incubated with 200 nM of RBD antigen in PBS for 1hr on ice. Cells were resuspended in 50µL of secondary stain containing streptavidin-PE (Invitrogen) at a 1:200 dilution and incubated for 30 min on ice. Cell binding was analyzed by flow cytometry using a Stratedigm S1300Exi Flow Cytometer equipped with a 96 well plate high throughput sampler. Rsulting data were analyzed using FlowJo (10.7.1).

### Probe Generation

SARS-CoV-2 RBD and ΔRBM constructs were expressed as dimeric murine-Fc (mFc; IgG1) fusion proteins containing a HRV 3C-cleavable C-terminal 8xHis and SBP tags and purified as described above. SBP-tagged RBD- and ΔRBM-mFc dimers were individually mixed with fluorescently labeled streptavidin, SA-BV650 and SA-BV786 (BioLegend), to form RBD-mFc-BV650 and ΔRBM-mFc-BV786 tetramers. SARS-CoV-2 spike with a C-terminal Strep II tag was labeled separately with StrepTactin PE and APC (IBA) to form spike-PE and -APC tetramers, respectively. Both labeling steps were performed for 30 min at 4 °C prior to sorting.

### Single B Cell Sorting

Naive B cells were purified from PBMCs using the MACS Human B Cell isolation kit (Miltenyi Biotec) and incubated with 25nM of each SARS-CoV-2 probe (RBD-mFc-BV650, ΔRBM-mFc-BV786, spike-PE, and spike-APC) for 30 min at 4°C. Cells were stained with anti-human CD19 (Alexa-700), CD3 (PerCP-Cy5), IgD (PE-Cy7), IgG (BV711), CD27 (BV510), LiveDead Violet (Invitrogen), and Calcien (Invitrogen) for an additional 30 min. RBM-specific naive B cells, defined as CD19^+^/CD3^-^/IgG^-^/IgD^+^/spike PE^+^/spike APC^+^/RBD^+^/ΔRBM^--^, were single-cell sorted using BD FACS Aria II (BD Biosciences) into 96-well plates containing lysis buffer supplemented with 1% BME. Within the CD19^+^/IgG^−^/IgD^+^ gated cells, we also confirmed that 97% of the events were CD27 negative. Plates were stored at -80 °C for subsequent analysis. Flow cytometry data was analyzed using FlowJo software version 10.7.1.

### BCR Sequencing

BCR Sequencing was carried out as described previously (*37*). Briefly, whole transcriptome amplification (WTA) was performed on the sorted cell-lysates according to the Smart-Seq2 protocol (*105*). We then amplified heavy and light chain sequences from the WTA products utilizing pools of partially degenerate pools of V region specific primers (Qiagen HotStar Taq Plus). Heavy and light chain amplifications were carried out separately, with each pool containing pooled primers against human IGHV and heavy chain constant region genes, or human IGLV, IGKV, and light chain constant region genes. Cellular barcodes and index adapters (based on Nextera XT Index Adapters, Illumina Inc.) were added using a step-out PCR method. Amplicons were then pooled and sequenced using a 250x250 paired end 8x8 index reads on an Illumina Miseq System. The data were then demultiplexed, heavy and light chain reads were paired, and overlapping sequence reads were obtained (Panda-Seq) (*106*) and aligned against the human IMGT database (*107*).

### Interferometry binding experiments

Interferometry experiments were performed using a BLItz instrument (ForteBio). Fabs (0.1 mg/ml) were immobilized on Ni-NTA biosensors. The SARS-CoV-2 RBD analyte was titrated (10µM, 5µM, 2.5µM, and 1µM) to acquire binding affinities; the *K_D_* was obtained through global fit of the titration curves by applying a 1:1 binding isotherm using vendor-supplied software.

### Pseudotyped neutralization assay

SARS-CoV-2 neutralization was assessed using lentiviral particles pseudotyped as previously described (*10, 108*). Briefly, lentiviral particles were produced via transient transfection of 293T cells. The titers of viral supernatants were determined via flow cytometry on 293T-ACE2 cells (*108*) and via the HIV-1 p24^CA^ antigen capture assay (Leidos Biomedical Research, Inc.). Assays were performed in 384-well plates (Grenier) using a Fluent Automated Workstation (Tecan). IgGs starting at 150 µg/ml, were serially diluted (3-fold) in 20µL followed by addition of 20 µL of pseudovirus containing 250 infectious units and incubated at room temperature for 1 hr. Finally, 10,000 293T-ACE2 cells (*108*) in 20 µL cell media containing 15 µg/ml polybrene were added to each well and incubated at 37 °C for 60-72 hrs. Following transduction, cells were lysed using a previously described assay buffer (*109*) and shaken for 5 min prior to quantitation of luciferase expression using a Spectramax L luminometer (Molecular Devices). Percent neutralization was determined by subtracting background luminescence measured from cells control wells (cells only) from sample wells and dividing by virus control wells (virus and cells only). Data were analyzed using Graphpad Prism.

### cryo-EM sample preparation, data collection and processing

SARS-CoV-2 spike HexaPro was incubated with ab090 Fab at 0.6 mg/mL at a molar ratio of 1.5:1 Fab:Spike for 20 minutes at 4°C and two 3 μl aliquots were applied to UltrAuFoil gold R0.6/1 grids and subsequently blotted for 3 seconds at blot force 3 twice, then plunge-frozen in liquid ethane using an FEI Vitrobot Mark IV. Grids were imaged on a Titan Krios microscope operated at 300 kV and equipped with a Gatan K3 Summit direct detector. 10,690 movies were collected in counting mode at 16e^−^/pix/s at a magnification of 81,000, corresponding to a calibrated pixel size of 1.058 Å. Defocus values were at around -2.00 μm. Micrographs were aligned and dose weighted using Relion’s (*110*) implementation of MotionCorr2 (*111*). Contrast transfer function estimation was done in GCTF (*112*). Particles were picked with crYOLO (*113*) with a model trained with 12 manually picked micrographs with particle diameter value of 330Å. Initial processing was performed in Relion. The picked particles were binned to ∼ 12Å/pixel and subjected to a 2D classification. Selected particles were then extracted to ∼6Å/pixel then subjected to a second round of 2D classification. An initial model was generated on the selected particles at ∼6Å/pixel and used as a reference for two rounds of 3D classification; first to select particles containing SARS-CoV-2 spike then to select particles containing both spike and ab090. Selected particles were unbinned then aligned using 3D auto-refine and subjected to a third round of 3D classification to select for a single class with SARS-CoV-2 spike bound with one ab090 Fab. Selected particles were aligned using 3D auto-refine before undergoing CTF refinement and Bayesian polishing. Polished particles were then simultaneously focus-aligned relative to the RBD and ab090 region (Figure S5 A) to aid in model building of this region of interest and imported to cryoSPARC (*114*). Imported particles were aligned using non-uniform refinement and local resolution estimation (Figure S5B). Non-uniform refined maps were then sharpened with DeepEMhancer then used to dock a previously built SARS-CoV-2-spike model (PDB ID 7LQW).

### cryo-EM model building

Backbone models were built by docking the variable regions of structurally similar Fabs (PDB ID 2D2P and 6FG1 for heavy and light chains, respectively) and a previously built RBD (6M0J) into the focus refined maps using UCSF Chimera (*115*) variable regions were then mutated and manually built using COOT (*116*). For the remainder of the spike, a previously published model (PDB ID 6VXX) was docked into the full, sharpened map in UCSF Chimera.

### RBD nanoparticle production and conjugation

Monomeric SARS-CoV-2 wild-type, B.1.1.7, B.1.351, and WIV1 RBDs were recombinantly produced and purified as described above with an 8xHis and SpyTag (cite) at the C-terminus. *Helicobacter pylori* ferritin nanoparticles (NP) were expressed separately with N-terminal 8xHis and SpyCatcher tags. SpyTag-SpyCatcher conjugations were performed overnight at 4°C with a 4-fold molar excess of SpyTag-RBD relative to SpyCatcher-NP. The conjugated RBD-NPs were repurified by size-exclusion chromatography to remove excess RBD-SpyTag.

### *In vitro* BCR triggering

The capacity of RBD-NPs to trigger naive BCR signaling was determined through activation of Ramos cells engineered to display mono-specific IgM BCRs of interest, as previously described (*76*). Briefly, BCRs for ab090 and ab072 were stably expressed in an IgM negative Ramos B cell clone by lentiviral transduction. Five to seven days post transduction, confluent BCR-expressing B cells were FACS sorted on IgM (APC anti-human IgM; BioLegend) and kappa light chain (PE anti-human kappa light chain; BD Biosciences) double positivity using a SH800S Cell Sorter (Sony Biotechnology). Sorted cells were expanded in RPMI (GIBCO) and evaluated for B cell activation by labeling 10 million cells with 0.5µg/ml Fura red dye (Invitrogen) in 2ml of RPMI at 37°C for 30 min. Cells were then washed and resuspended to 4 million cells/ml in RPMI. BCR triggering was measured in response to the RBD-NPs described above by flow cytometry (LSR II, BD Biosciences) as the ratio of Ca^2+^ bound/unbound states of Fura red. Ratiometric measures for individual B cell lines were normalized to the maximum Ca^2+^ flux as measured by exposure to 10µg/ml ionomycin.

### *in vitro* affinity maturation of ab090 and ab072

To build yeast display libraries for ab090 and ab072, variable heavy and light chains were reformatted into an scFv and synthesized as gBlocks (IDT). gBlocks were amplified by polymerase chain reaction (PCR) using Q5 polymerase (New England BioLabs) following the manufacturer’s protocol. The amplified DNA purified and subsequently mutagenized by error-prone PCR (ePCR) via the GeneMorph II Random Mutagenesis Kit (Agilent Technologies) with a target nucleotide mutation frequency of 0-4.5 mutations/kb. Mutagenized scFv DNA products were combined with the linearized yeast display vector pCHA (*117*) and electroporated into EBY100 grown to mid-log phase in YPD media, where the full plasmid was reassembled by homologous recombination (*85*). The final library size was estimated to be 4 x 10^7^.

The scFv libraries and selection outputs were passaged in selective SDCAA media (20 g/L dextrose, 6.7 g/L Yeast Nitrogen base, 5 g/L Bacto casamino acids, 5.4 g/: Na_2_HPO_4_ and 8.56 g/L NaH_2_PO_4_·H_2_O) at shaking at 30°C and induced in SGCAA media (same as SDCAA wit 20 g/L galactose instead of dextrose) at 20°C. The scFv libraries were induced covering at least 10-fold of their respective diversities and subject to three rounds of selection for binding to SBP-tagged SARS-CoV-2 RBD-Fc. Induced yeast libraries were stained for antigen binding (RBD-Fc APC tetramers) and scFv expression (chicken anti-c-myc IgY; Invitrogen). Following two washes in PBSF (1x PBS, 0.1% w/v BSA), yeast was stained with donkey anti-chicken IgY AF488 (Jackson ImmunoResearch). Two gates were drawn for cells with improved RBD binding over parental clones, a more stringent “edge” gate represented ∼1% and a “diversity” gate represented ∼3-5% of the improved output. Yeast from the final round of selection were resuspended in SDCAA media and plated on SDCAA agar plates for single colony isolation and Sanger sequencing from which IgGs and Fabs were cloned and recombinantly expressed as described above.

## Acknowledgements

We thank members of the Schmidt and Lingwood Labs for helpful discussions, especially Tim Caradonna, Catherine Jacob-Dolan, and Daniel Maurer. We thank Samuel Kazer, James Gatter and Alex Shalek for BCR sequencing advice, Jason McLellan for the SARS-CoV-2 spike plasmid, and Nir Hacohen and Michael Farzan for ACE2 expressing 293T cells. Some of this work was performed at the National Center for CryoEM Access and Training (NCCAT) and the Simons Electron Microscopy Center located at the New York Structural Biology Center, supported by the NIH Common Fund Transformative High Resolution Cryo-Electron Microscopy program (U24 GM129539), and by grants from the Simons Foundation (SF349247) and NY State Assembly.

## Author contributions

J.F., J.B., A.B.B., G.B., D.L., A.G.S. designed research; J.F., J.B., C.G.A., K.S.D., E.C.L, B.M.H., L.R., M.S., T.B.M., V.O., N.H., performed research; J.F., J.B., C.G.A, G.B., D.L., A.G.S analyzed data; J.F. and A.G.S. wrote the paper. J.F., J.B., C.G.A., B.M.H, A.B.B., G.B., D.L., A.G.S edited and commented on the paper.

## Data and materials availability

All data are provided in the Supplementary Materials. Requests for material should be addressed to Daniel Lingwood or Aaron G. Schmidt. This work is licensed under a Creative Commons Attribution 4.0 International (CC BY 4.0) license, which permits unrestricted use, distribution, and reproduction in any medium, provided the original work is properly cited. To view a copy of this license, visit https://creativecommons.org/licenses/by/4.0/. This license does not apply to figures/photos/artwork or other content included in the article that is credited to a third party; obtain authorization from the rights holder before using such material. The EM maps have been deposited in the Electron Microscopy Data Bank (EMDB) under accession code: EMD-24279.

## Supplemental Information

**fig. S1.**
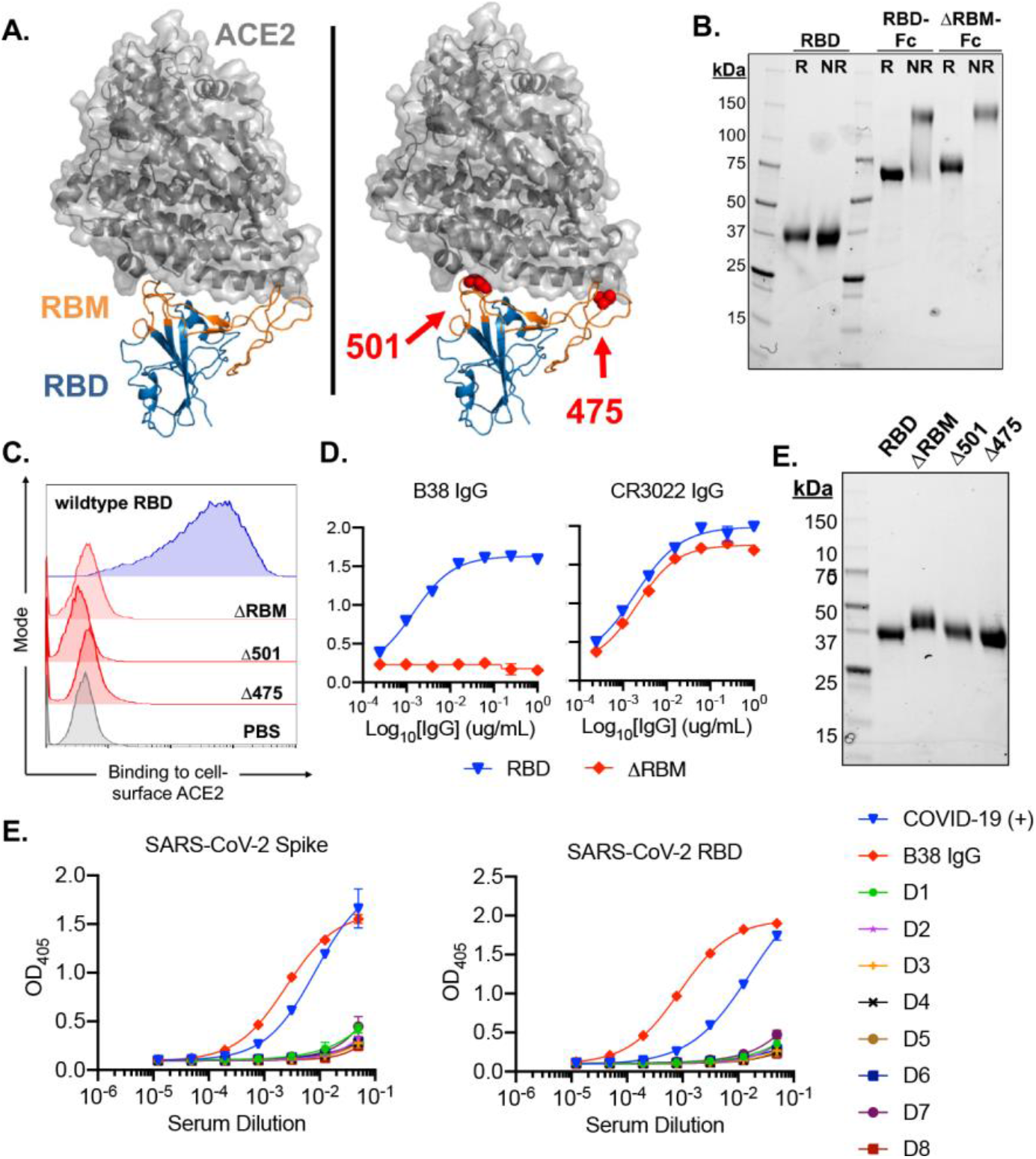
Design and characterization of SARS-CoV-2 antigens and healthy donor sera binding. (**A**) SARS-CoV-2 RBD in complex with viral receptor, ACE2 shown in blue and grey, respectively (PDB 6M0J). Wild-type RBD with, the receptor binding motif (RBM), shown in orange (left panel). Structural model of the ΔRBM probe designed to abrogate binding to ACE2 (right panel). Putative N-linked glycosylation sites engineered onto the RBM are shown in red spheres at amino acid positions 501 and 475. (**B**) SDS-PAGE gel under reducing (R) and non-reducing (NR) conditions for monomeric RBD, RBD-Fc and ΔRBM-Fc. (**C**) Wildtype RBD, ΔRBM and single glycan variant binding to ACE2-expressing 293T cells by flow cytometry. Wild-type RBD binding shown in blue, glycan variant binding shown in red. Streptavidin-PE was used to detect the relative intensity of antigen binding to cell-surface ACE2. A PBS control (gray) indicates secondary-only staining. (**D**) Control antibody ELISA binding to RBD and ΔRBM antigens. RBM-specific antibody, B38 (left). Non-RBM-specific control antibody, CR3022 (right). (**E**) ΔRBM and Δ501 and Δ475 variants analyzed by SDS-PAGE gel under reducing conditions; wildtype RBD is shown for comparison. (E) SARS-CoV-2 spike (left) and RBD (right) sera ELISA from human subjects 1-8. Sera from a COVID-19 convalescent patient and control antibody, B38, were included as positive controls.

**fig. S2.**
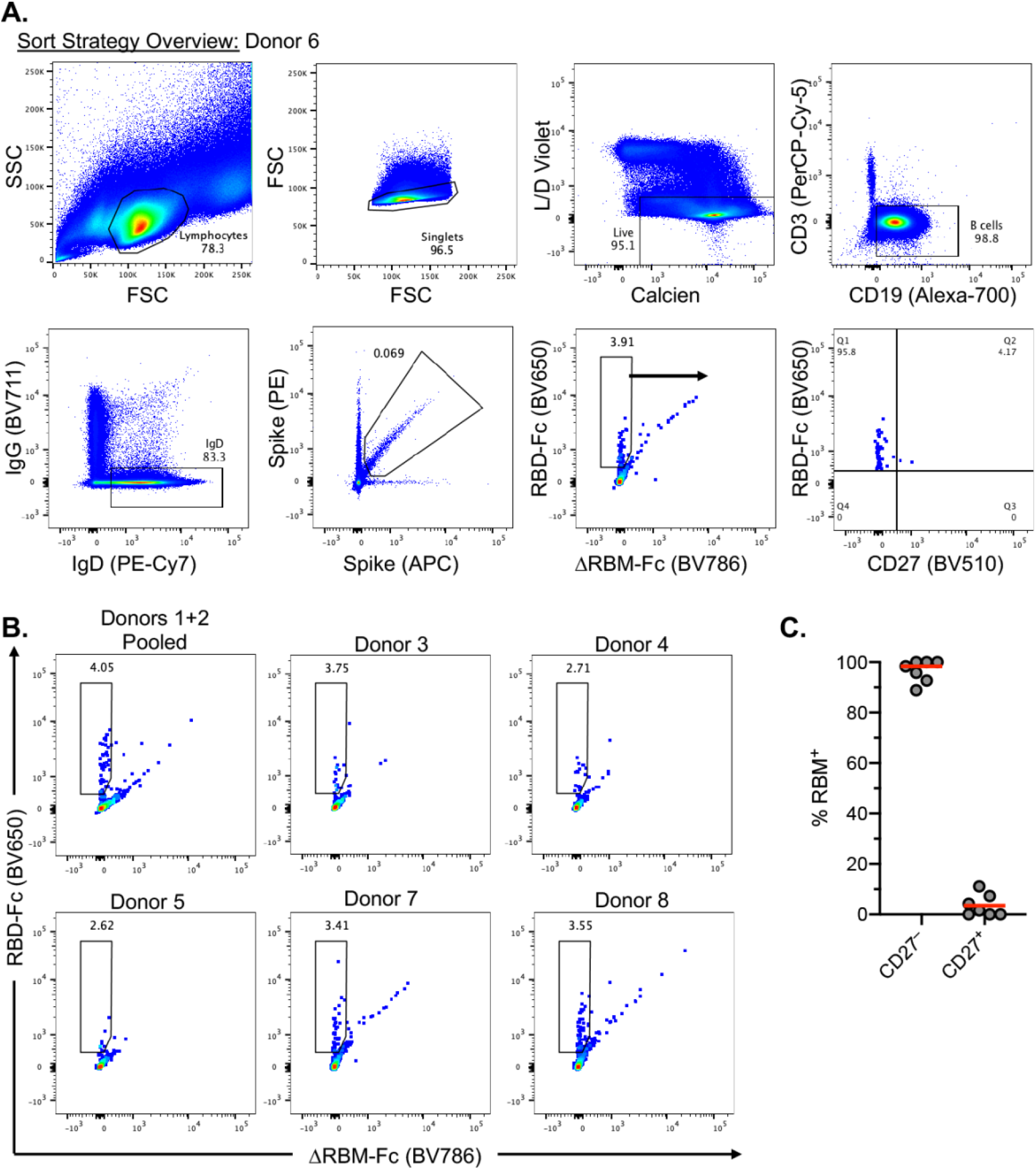
PBMC flow cytometry analyses. (**A**) Representative gating strategy used for FACS of PBMCs pooled from donors 1 and 2. Gating was on naive B cells defined by single living lymphocytes that were CD19^+^CD3^-^IgD^+^IgG^-^. Sorted cells were RBM-specific as defined by spike-PE^+^/spike-APC^+^/RBD-Fc-BV650^+^/ΔRBM-Fc-BC650-. Sort gate is denoted by the blue arrow. The bottom right plot shows CD27 staining of sorted RBM-specific naive B cells. (**B**) Flow cytometry showing the sort gate and percentage of RBM-specific B cells for the remaining 6 healthy human donors. (**C**) RBM-specific B cell frequency among CD27^+^ and CD27^-^ cells. Each symbol represents a different donor (*n* = 8).

**fig. S3.**
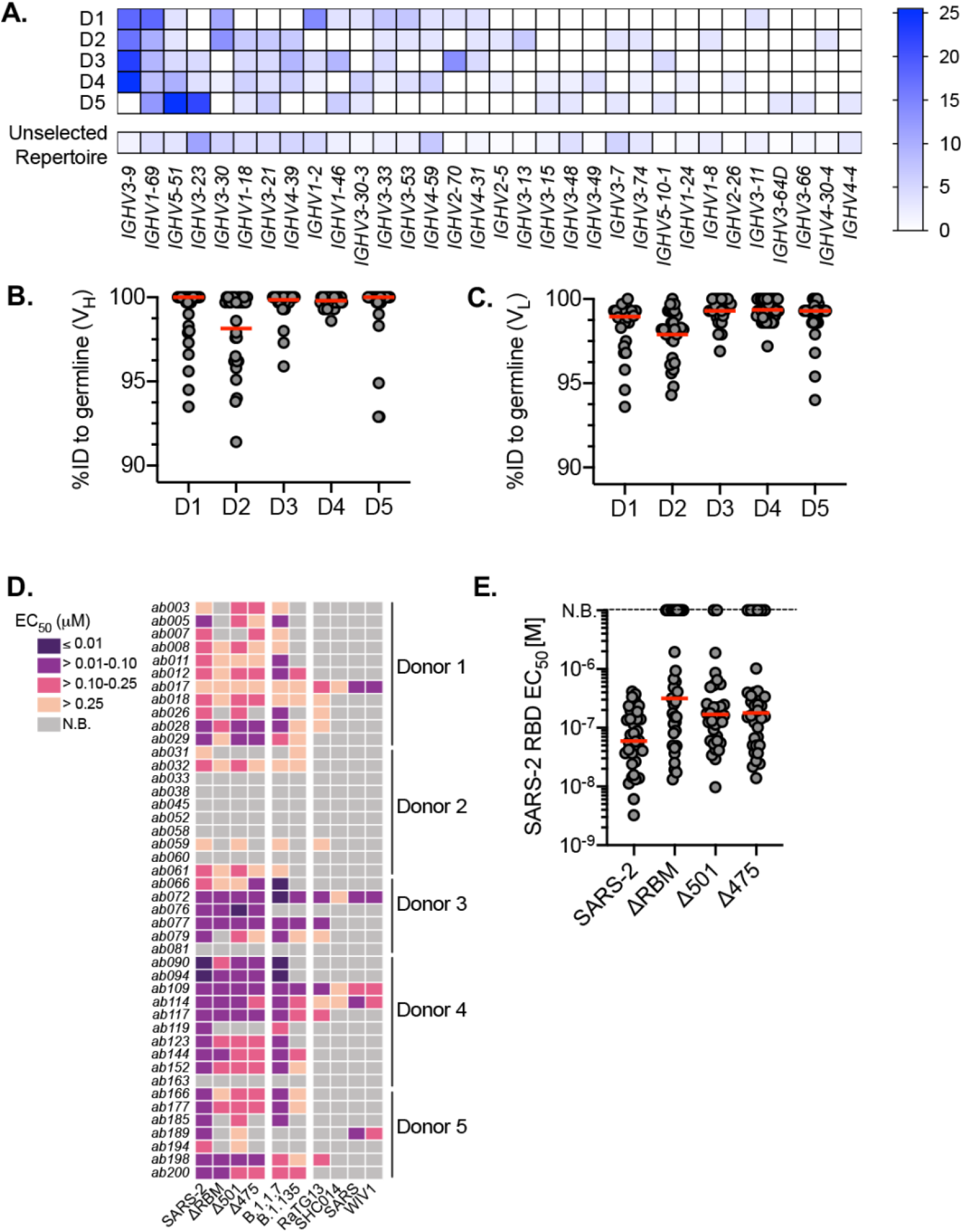
Repertoire comparison, germline identity, and IgG binding by individual donor. (**A**) Heatmap showing V_H_-gene usage of isolated antibodies derived from donors 1-5. Unselected repertoire gene usage derive from a high-throughput sequencing data set of circulating B cells across 10 human subjects (*46*). Heatmap scale represents percent of total paired sequence from each donor. Divergence from inferred germline gene sequences separated by individual donor for (**B**) V_H_ and (**C**) V_L_. Red bars indicate the median percent values, and each dot represents an individual paired sequence. (**D**) Heatmap showing IgG binding to RBDs (*n* = 44) sorted by donor. **(E**) ELISA EC_50_ values for IgGs with detectable SARS-CoV-2 RBD binding (*n* = 36) against RBM glycan probes. Red bars indicate the mean EC_50_ values.

**fig. S4.**
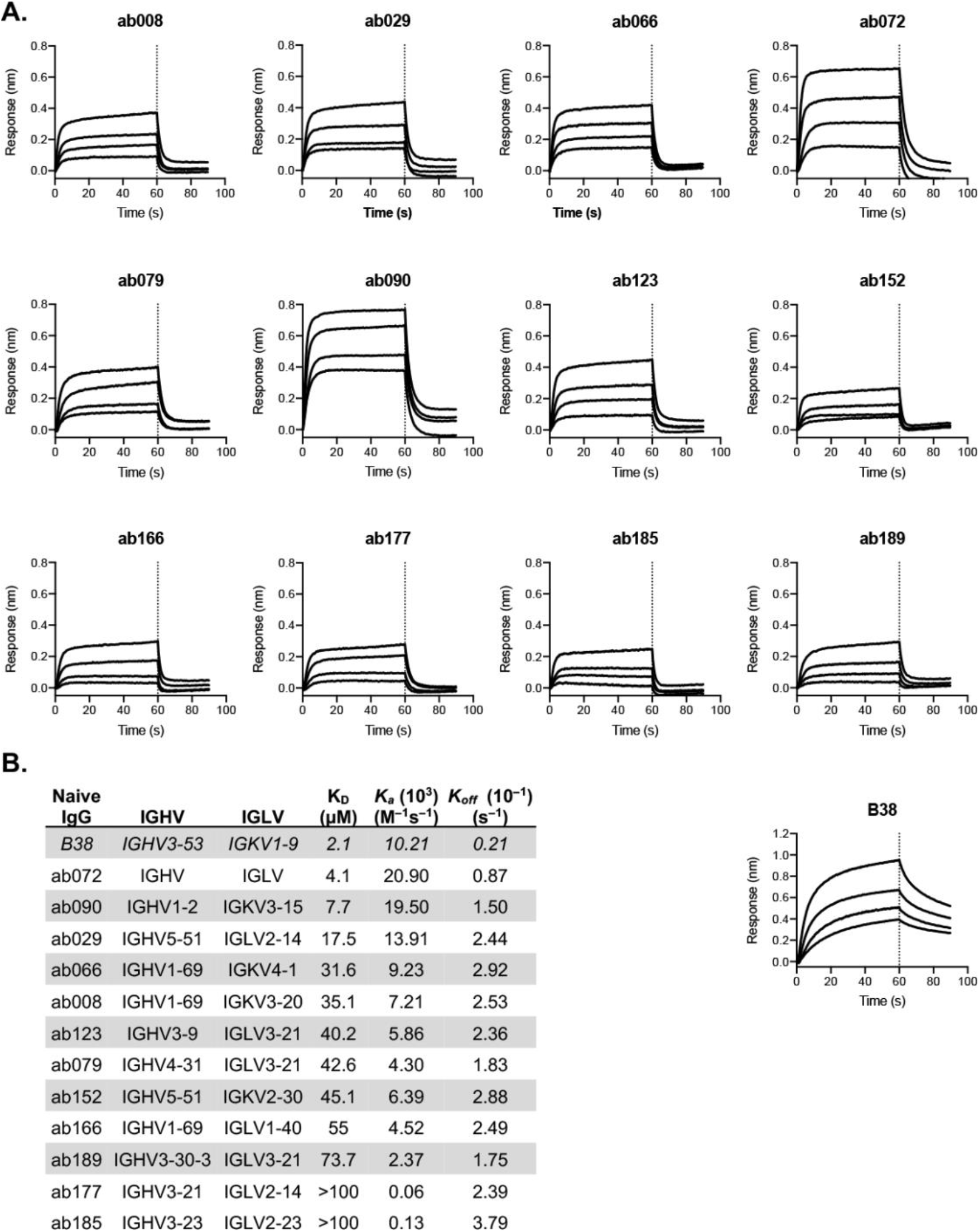
SARS-CoV-2 RBD-binding kinetics of isolated naive antibodies. (**A**) Biolayer interferometry (BLI) binding kinetic analysis of titrated SARS-CoV-2 RBD to immobilized Fabs. Dotted line at 60 s denotes the start of the dissociation phase. (**B**) Kinetic and equilibrium constants for binding to RBD calculated from a 1:1 binding model using a global fit to all curves for each Fab using vendor supplied software. B38 Fab is used as a positive control.

**fig. S5.**
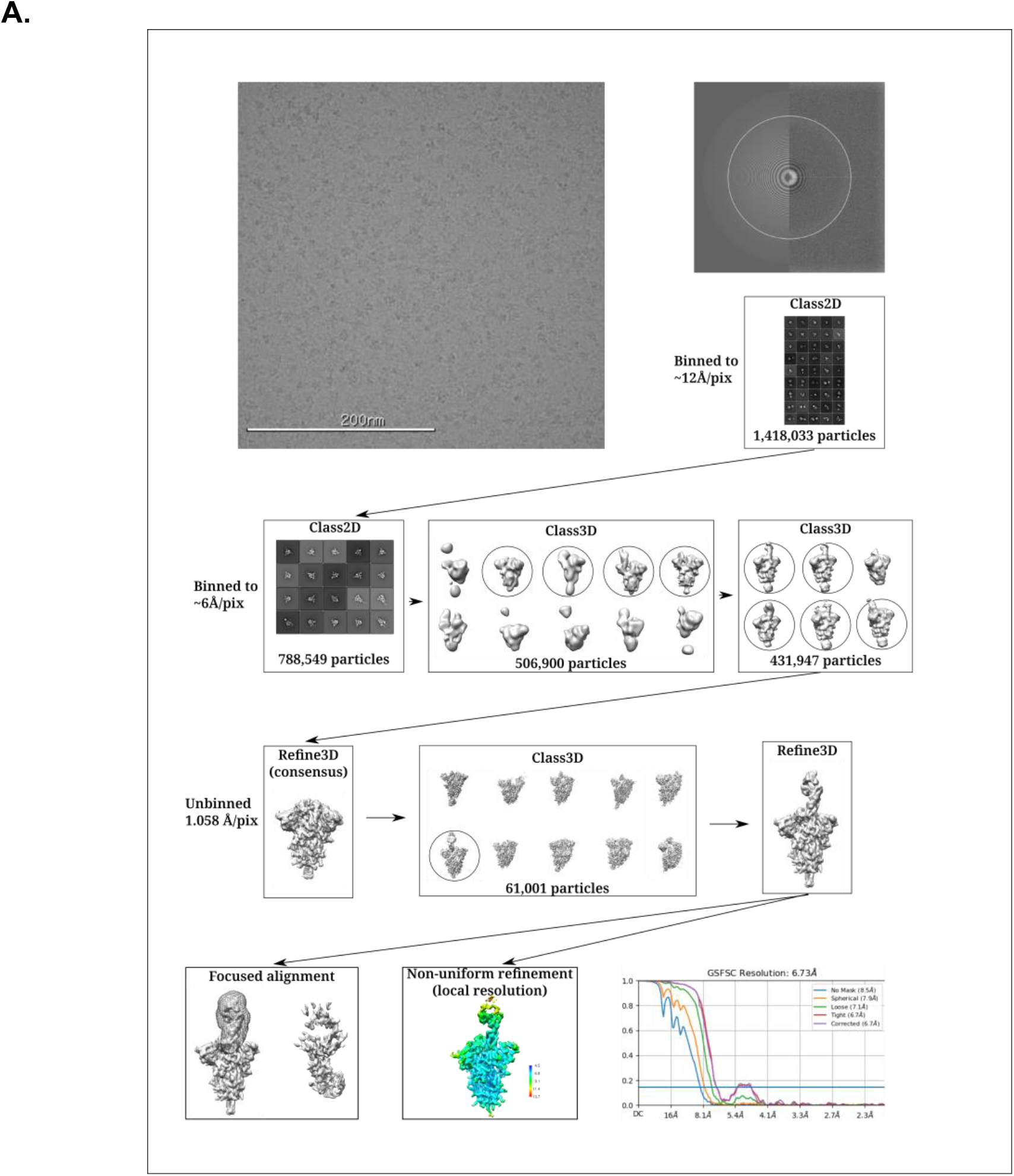
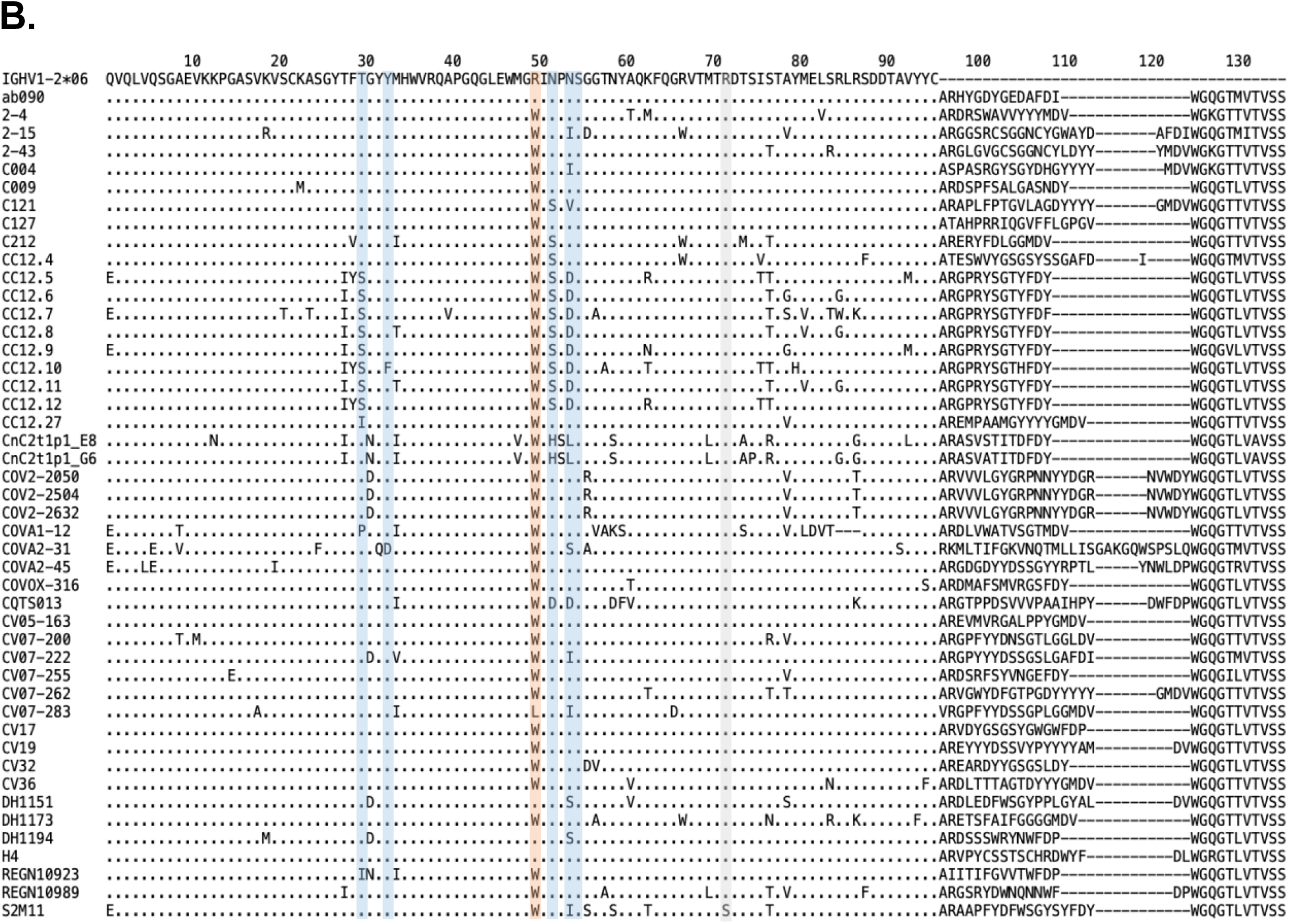
Structural characterization and analysis. **(A)** Cryo-EM data processing scheme of ab090 Fab bound with SARS-CoV-2 spike. See the Methods section for more details. (**B**) Heavy chain amino acid sequence alignment of ab090 with IGHV1-2 derived antibodies from convalescent COVID-19 patients. Sequences were obtained from CoV-AbDab (*118*) and aligned to the IGHV1-2*06 reference. Residues forming the germline-encoded HCDR1 and HCDR2 motif contacting the SARS-CoV-2 RBD are highlighted in blue. The single nucleotide polymorphism in the *06 allele at position 50 is highlighted red. The site of the dominant mutation from *in vitro* affinity maturation efforts with ab090 is highlighted in green.

**fig. S6.**
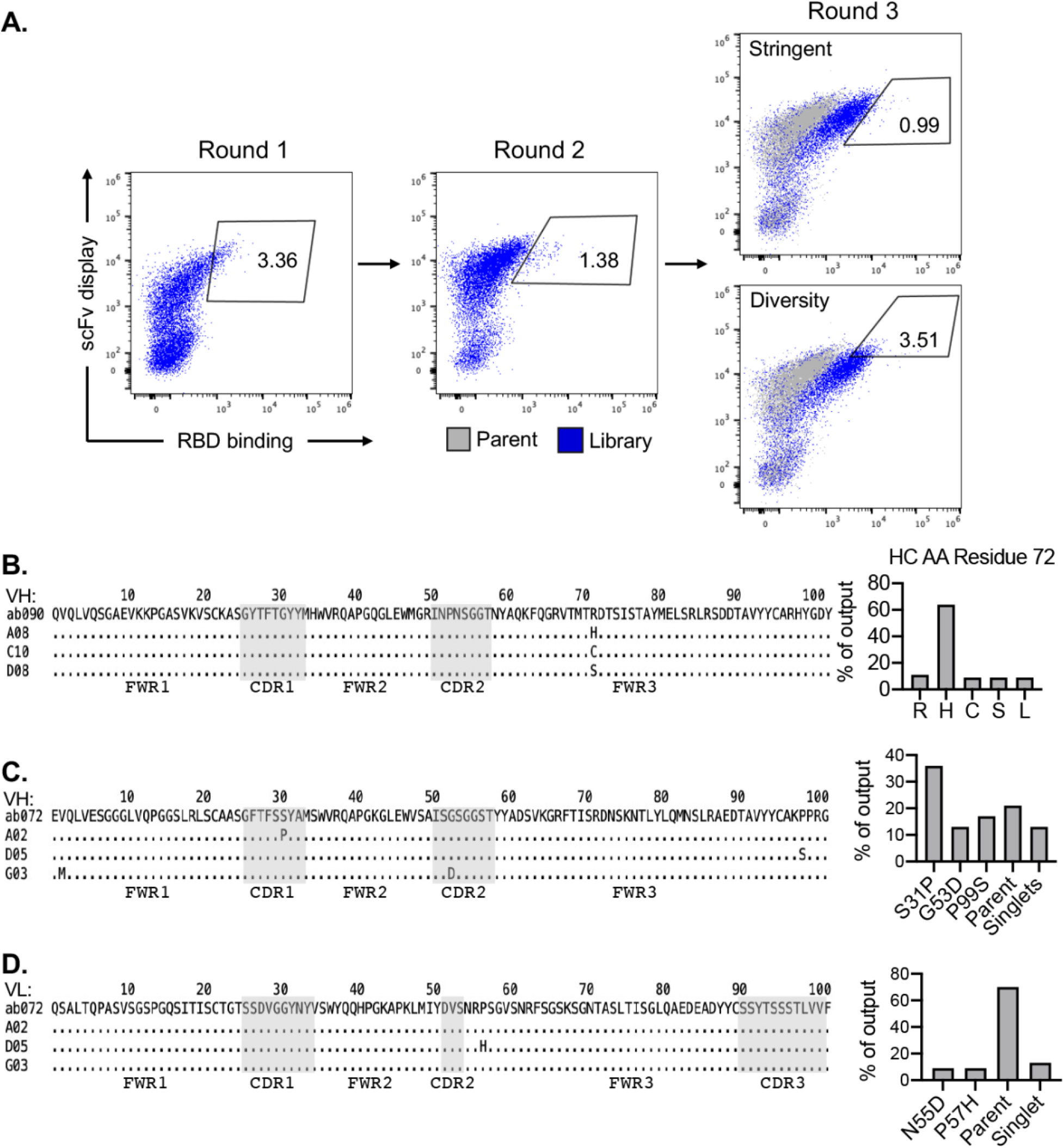
Representative affinity maturation selection strategy and output sequence overview. (**A**) Flow cytometric sorting of diversified single chain variable fragment (scFv) libraries of ab090. Gates represent the yeast population sorted for subsequent selections. After 2 rounds of enrichment for wildtype SARS-CoV-2 binding, a “stringent” and “diversity gate were sorted in round 3 indicating the yeast populations sorted for individual colony isolation and sequencing. Alignment of the V_H_ sequencing output clones for ab090 (**B**) and ab072 (**C**) with the output frequency of each mutation from a total of 48 single colonies. (**D**) Alignment of the V_L_ sequencing output clones ab072 with the output frequency of each mutation from a total of 48 single colonies. The V_L_ output for ab090 was exclusively parent.

